## Supplementary Information combined for "Intrinsic multiplication rate variation and plasticity of human blood stage malaria parasites"

**Supplementary Table S1.** European Nucleotide Archive accession numbers for Illumina paired-end short-read whole genome sequence data for each of the *P. falciparum* clinical isolates sampled after 25, 77 and 153 days of continuous culture.

| Patient Isolate | European Nucleotide Archive accession numbers: | | |
|---|---|---|---|
| | Day 25 | Day 77 | Day 153 |
| **271** | ERR2234984 | ERR2496504 | ERR2234997 |
| **272** | ERR2234970 | ERR2508979 | ERR2508991 |
| **273** | ERR2234999 | ERR2508980 | ERR2496509 |
| **274** | ERR2235001 | ERR2508981 | ERR2496510 |
| **275** | ERR2234972 | ERR2234974 | ERR2496511 |
| **276** | ERR2436108, 2436120, 2436132 | ERR2508982 | ERR2496512 |
| **277** | ERR2508983 | ERR2234976 | - |
| **278** | ERR2235003 | ERR2508984 | ERR2234986 |
| **279** | ERR2234978 | ERR2235005 | ERR2508992 |
| **280** | ERR2234980 | ERR2234982 | ERR2234987 |
| **281** | ERR2234985 | ERR2234971 | - |
| **282** | ERR2234973 | ERR2234975 | ERR2508993 |
| **284** | ERR2436109, 2436121, 2436133 | ERR2234977 | ERR2496515 |
| **285** | ERR2235006 | ERR2235007 | ERR2496516 |
| **286** | ERR2508986 | ERR2436110, 2436122, 2436134 | ERR2234989 |
| **287** | ERR2235008 | ERR2234979 | ERR2508994 |
| **288** | ERR2436111, 2436123, 2436135 | ERR2508987 | ERR2496517 |
| **289** | ERR2508988 | ERR2508989 | - |
| **290** | ERR2436112, 2436124, 2436136 | ERR2234981 | ERR2496519 |
| **291** | ERR2235009 | ERR2235000 | ERR2508995 |
| **292** | ERR2234983 | ERR2496506 | ERR2496520 |
| **293** | ERR2235002 | ERR2235004 | ERR2508996 |
| **294** | - | ERR2496507 | ERR2508997 |
| **296** | ERR2436114,2436126,2436138 | ERR2496508 | - |

**Supplementary Table S2.** Genome-wide sequence coverage (percentage of genome covered with > 5 reads) and within-isolate fixation indices ($F_{WS}$) for each of the *P. falciparum* clinical isolates sampled after 25, 77 and 153 days of continuous culture.

| Patient Isolate | Genome-wide percent coverage: | | | $F_{WS}$ index: | | |
|---|---|---|---|---|---|---|
| | **Day 25** | **Day 77** | **Day 153** | **Day 25** | **Day 77** | **Day 153** |
| **271** | 86.7 | 53.9 | 88.7 | 0.987 | 0.992 | 0.987 |
| **272** | 89.4 | 91.1 | 91.1 | 0.975 | 0.985 | 0.972 |
| **273** | 89.4 | 90.9 | 65.2 | 0.812 | 0.984 | 0.990 |
| **274** | 89.3 | 91.9 | 52.6 | 0.304 | 0.472 | 0.995 |
| **275** | 88.7 | 86.9 | 63.1 | 0.762 | 0.829 | 0.990 |
| **276** | 91.3 | 91.4 | 61.1 | 0.748 | 0.880 | 0.857 |
| **277** | 91.1 | 87.4 | - | 0.617 | 0.976 | - |
| **278** | 89.0 | 91.0 | 89.6 | 0.534 | 0.975 | 0.990 |
| **279** | 88.9 | 88.0 | 89.9 | 0.837 | 0.544 | 0.916 |
| **280** | 88.0 | 88.7 | 88.2 | 0.971 | 0.957 | 0.976 |
| **281** | 88.5 | 88.5 | - | 0.278 | 0.912 | - |
| **282** | 88.3 | 88.4 | 91.3 | 0.364 | 0.789 | 0.661 |
| **284** | 90.7 | 87.0 | 68.7 | 0.727 | 0.516 | 0.957 |
| **285** | 89.6 | 89.9 | 52.6 | 0.382 | 0.607 | 0.984 |
| **286** | 91.1 | 90.8 | 90.0 | 0.639 | 0.940 | 0.867 |
| **287** | 88.6 | 88.9 | 90.3 | 0.985 | 0.989 | 0.980 |
| **288** | 91.3 | 90.7 | 65.3 | 0.293 | 0.710 | 0.990 |
| **289** | 91.0 | 90.9 | - | 0.985 | 0.979 | - |
| **290** | 91.6 | 88.6 | 57.6 | 0.413 | 0.884 | 0.987 |
| **291** | 89.8 | 88.9 | 90.0 | 0.477 | 0.978 | 0.951 |
| **292** | 88.2 | 51.1 | 62.8 | 0.986 | 0.994 | 0.990 |
| **293** | 89.2 | 89.4 | 90.9 | 0.961 | 0.989 | 0.973 |
| **294** | - | 89.9 | 92.0 | - | 0.237 | 0.338 |
| **296** | 90.7 | 53.4 | - | 0.991 | 0.991 | - |

The genome-wide SNP and indel calls for all individual samples at all time points are given at ftp://ngs.sanger.ac.uk/production/malaria/Resource/28.

**Supplementary Table S3.** Gametocyte counts as a proportion of all parasite stages in the cultured *P. falciparum* clinical isolates

| Patient Isolate | % Gametocyte counts in maintenance culture | | |
|---|---|---|---|
| | Day 25 | Day 77 | Day 153 |
| **271** | 5 (60) | 4 (332) | 2 (125) |
| **272** | 6 (52) | 1 (409) | 0 (234) |
| **273** | 7 (110) | 1 (276) | 1 (220) |
| **274** | 5 (100) | 3 (289) | 0 (397) |
| **275** | 5 (76) | 2 (311) | 1 (128) |
| **276** | 1 (86) | 3 (392) | 0 (318) |
| **277** | 1 (82) | 1 (254) | |
| **278** | | 1 (336) | 6 (157) |
| **279** | 7 (61) | 2 (280) | |
| **280** | 3 (89) | 2 (439) | 2 (202) |
| **281** | | 2 (311) | |
| **282** | 10 (42) | 2 (292) | 0 (245) |
| **283** | 0 (87) | 3 (279) | 0 (285) |
| **284** | 9 (85) | 4 (112) | 2 (130) |
| **285** | 7 (60) | 3 (102) | 1 (284) |
| **286** | 2 (139) | 2 (312) | 1 (193) |
| **287** | | 2 (124) | 0 (108) |
| **288** | | 0 (200) | 0 (338) |
| **289** | 1 (89) | 1 (146) | 1 (360) |
| **290** | | | 2 (301) |
| **291** | 10 (69) | 0 (324) | 1 (446) |
| **292** | 2 (116) | 0 (296) | 1 (180) |
| **293** | 9 (89) | 2 (283) | 0 (389) |
| **294** | 5 (22) | 3 (265) | 0 (205) |

Percentages of gametocytes are presented from parasite stage-differential counts made on the day and up to 7 days before or after when parasites were taken for each of the three timepoint assays. Numbers of parasites counted are shown in brackets. Blank cells correspond to points at which stage-differential counts were not performed on a given isolate.

**Supplementary Table S4. Multiple regression correlation of patient variables with parasite multiplication rates assayed after different lengths of time of culture.**

| | Day 25 culture | | Day 77 culture | | Day 153 culture | |
|---|---|---|---|---|---|---|
| | Coefficient | P value | Coefficient | P value | Coefficient | P value |
| Age | 1.43 | 0.17 | 2.35 | 0.03* | 1.48 | 0.15 |
| Haemoglobin level | -1.57 | 0.13 | -0.62 | 0.54 | -0.001 | 0.99 |
| Parasitaemia | 2.85 | 0.013* | 3.43 | 0.003* | 3.03 | 0.008* |

Adjusting for covariates, the residual coefficients tabulated are equivalent to z-scores, with P values indicating probability of these values overlapping with zero (null expectations). Asterisks indicate significant values.

**Supplementary Fig. S1.** Exponential growth assay data plotted for each of the individual clinical isolates tested in triplicate in erythrocytes from three different donors. The genome copy numbers measured by qPCR are plotted at each of the assay timepoints for each of the triplicates. Squared correlation coefficients of the assay timepoints including all replicates are shown, with all included assays having $r^2$ values above 0.90. The estimated multiplication rates (MR) per 48 hours are shown (95% confidence intervals of the estimates are given in brackets). **A.** Assays after 25 days of culture (18 isolates assayed successfully). **B.** Assays after 77 days of culture (23 isolates assayed successfully). **C.** Assays after 153 days of culture (19 isolates assayed successfully).

**A Day 25**

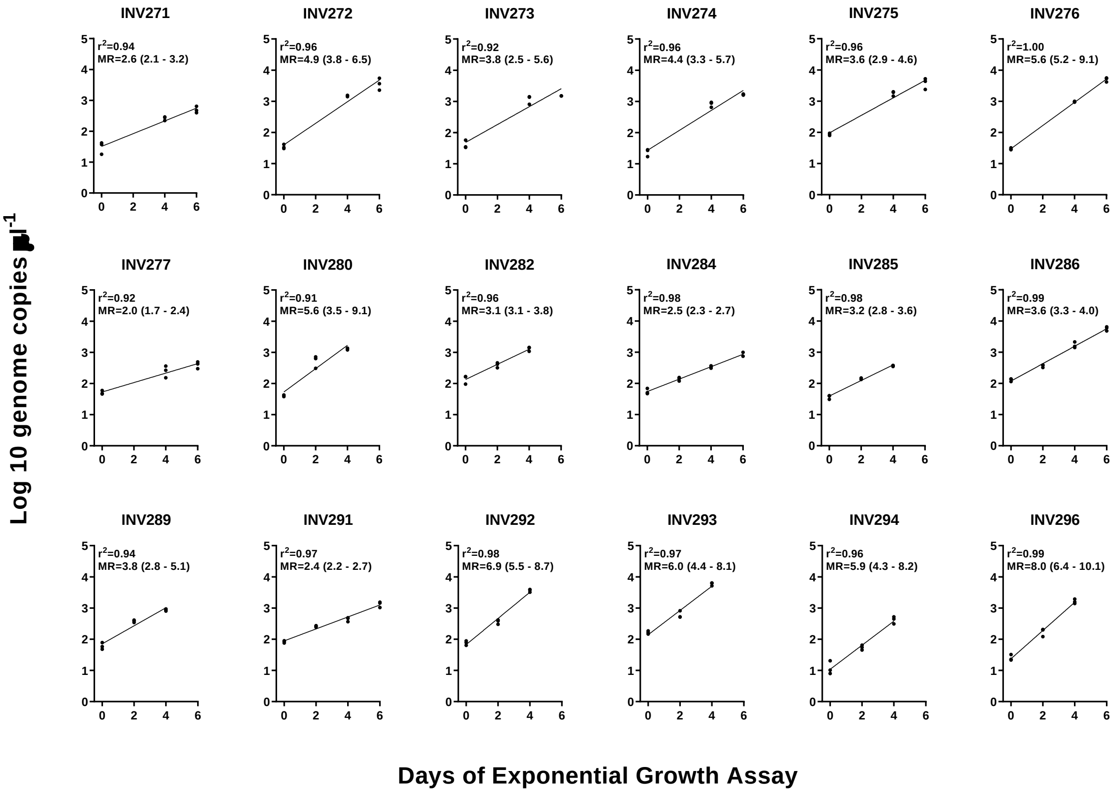

**B Day 77**

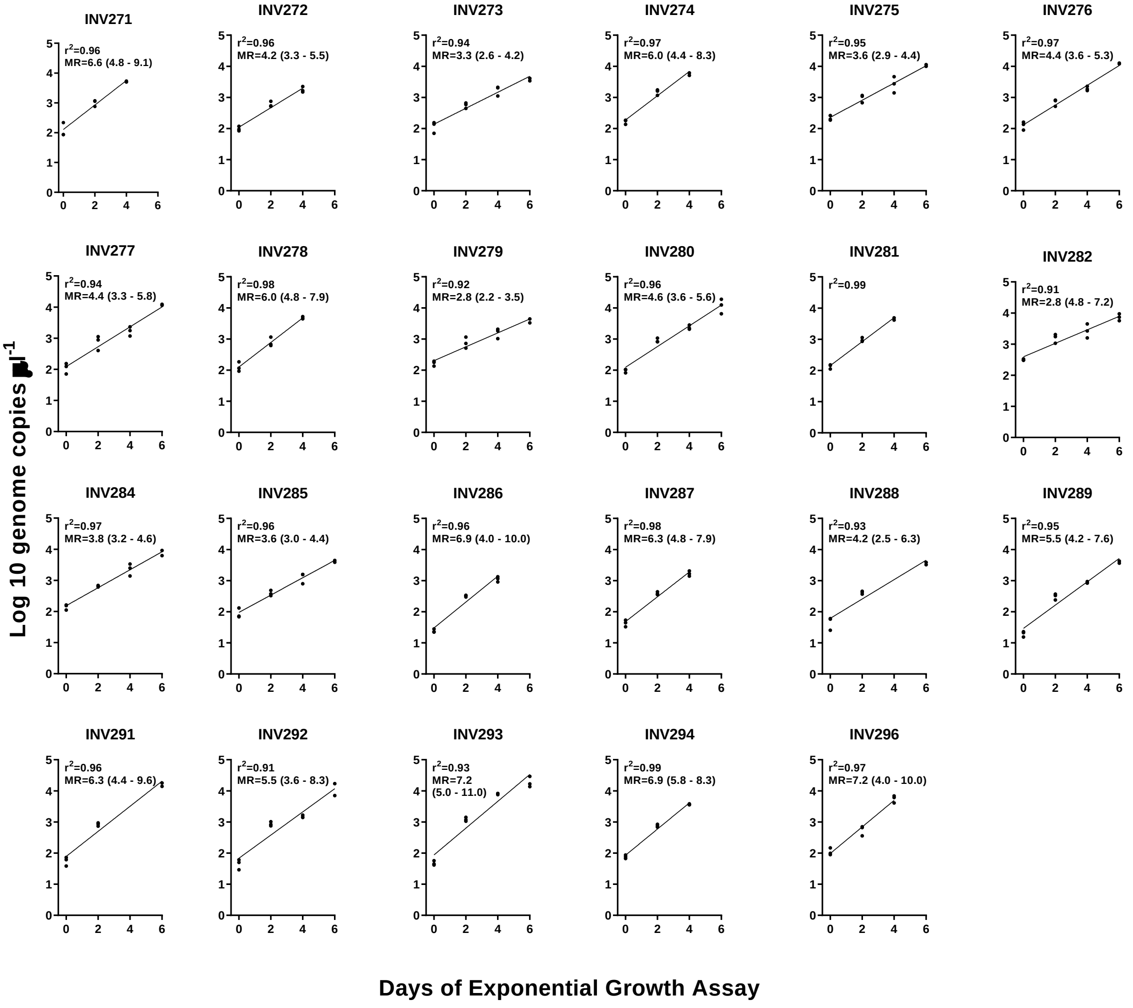
INV271
r²=0.96
MR=6.6 (4.8 - 9.1)
INV272
r²=0.96
MR=4.2 (3.3 - 5.5)
INV273
r²=0.94
MR=3.3 (2.6 - 4.2)
INV274
r²=0.97
MR=6.0 (4.4 - 8.3)
INV275
r²=0.95
MR=3.6 (2.9 - 4.4)
INV276
r²=0.97
MR=4.4 (3.6 - 5.3)
INV277
r²=0.94
MR=4.4 (3.3 - 5.8)
INV278
r²=0.98
MR=6.0 (4.8 - 7.9)
INV279
r²=0.92
MR=2.8 (2.2 - 3.5)
INV280
r²=0.96
MR=4.6 (3.6 - 5.6)
INV281
r²=0.99
INV282
r²=0.91
MR=2.8 (4.8 - 7.2)
INV284
r²=0.97
MR=3.8 (3.2 - 4.6)
INV285
r²=0.96
MR=3.6 (3.0 - 4.4)
INV286
r²=0.96
MR=6.9 (4.0 - 10.0)
INV287
r²=0.98
MR=6.3 (4.8 - 7.9)
INV288
r²=0.93
MR=4.2 (2.5 - 6.3)
INV289
r²=0.95
MR=5.5 (4.2 - 7.6)
INV291
r²=0.96
MR=6.3 (4.4 - 9.6)
INV292
r²=0.91
MR=5.5 (3.6 - 8.3)
INV293
r²=0.93
MR=7.2
(5.0 - 11.0)
INV294
r²=0.99
MR=6.9 (5.8 - 8.3)
INV296
r²=0.97
MR=7.2 (4.0 - 10.0)
Log 10 genome copies µl⁻¹
Days of Exponential Growth Assay

**C Day153**

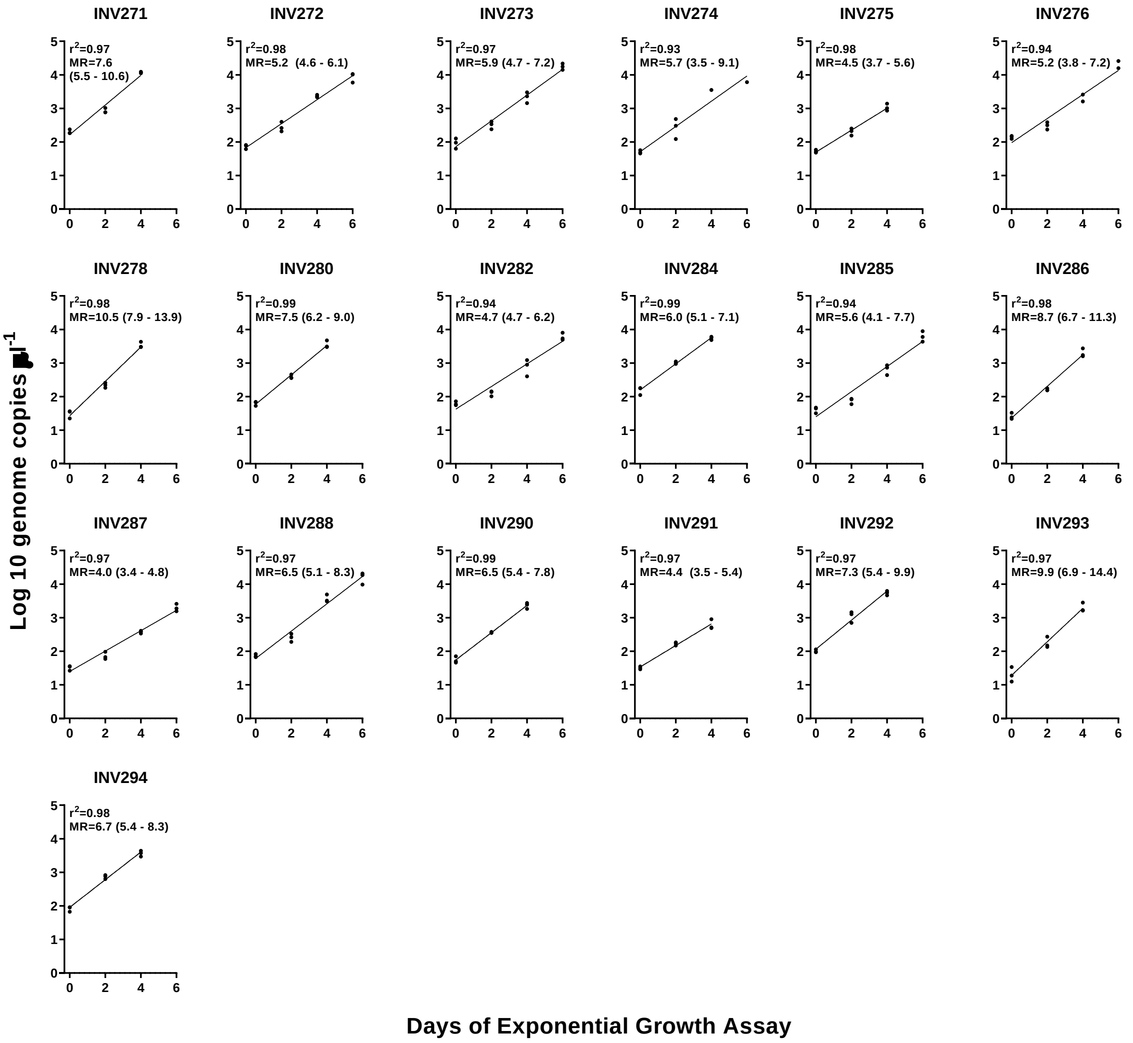
INV271
r2=0.97
MR=7.6
(5.5 - 10.6)
INV272
r2=0.98
MR=5.2 (4.6 - 6.1)
INV273
r2=0.97
MR=5.9 (4.7 - 7.2)
INV274
r2=0.93
MR=5.7 (3.5 - 9.1)
INV275
r2=0.98
MR=4.5 (3.7 - 5.6)
INV276
r2=0.94
MR=5.2 (3.8 - 7.2)
INV278
r2=0.98
MR=10.5 (7.9 - 13.9)
INV280
r2=0.99
MR=7.5 (6.2 - 9.0)
INV282
r2=0.94
MR=4.7 (4.7 - 6.2)
INV284
r2=0.99
MR=6.0 (5.1 - 7.1)
INV285
r2=0.94
MR=5.6 (4.1 - 7.7)
INV286
r2=0.98
MR=8.7 (6.7 - 11.3)
INV287
r2=0.97
MR=4.0 (3.4 - 4.8)
INV288
r2=0.97
MR=6.5 (5.1 - 8.3)
INV290
r2=0.99
MR=6.5 (5.4 - 7.8)
INV291
r2=0.97
MR=4.4 (3.5 - 5.4)
INV292
r2=0.97
MR=7.3 (5.4 - 9.9)
INV293
r2=0.97
MR=9.9 (6.9 - 14.4)
INV294
r2=0.98
MR=6.7 (5.4 - 8.3)
Log 10 genome copies µl-1
Days of Exponential Growth Assay

**Supplementary Fig. S2.** Comparison of multiplication rates in *P. falciparum* clinical isolates containing single genome sequences ($F_{WS}$ > 0.95) or mixed genome sequences ($F_{WS}$ < 0.95) at each of the timepoints of testing in culture. Each of the panels shows the isolate status in different ways, and numbers of isolates with multiplication rate assay and sequence data at each timepoint varies (all data on individual isolates are given in Table 1 and Supplementary Table 2). **A.** Each point represents an individual isolate, with 'days in culture' referring to the multiplication rate assay data only, so that black points represent isolates that had single genome sequences at *all* timepoints examined, white points represent those that had mixed genome sequences at *any* timepoint. **B.** Each point represents an individual isolate, shading of points representing whether they had single genome sequences (black) or mixed genome sequences (white) at each individual timepoint separately.

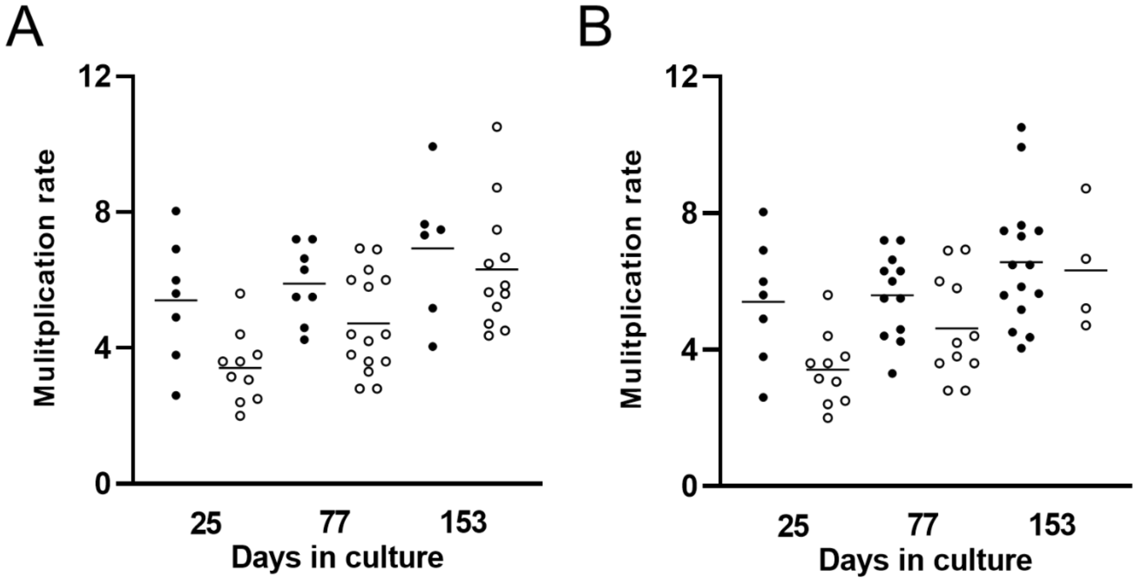

**Supplementary Fig. S3.** Numbers of merozoites per mature schizont counted in each of nine cultured clinical isolates and a test for correlation with multiplication rates at the final assayed timepoint. Counts of merozoites per mature schizont (with chemical blocking of egress using E64) were performed after different lengths of time in culture (closer to the middle or final assayed timepoints than to the first timepoint). **A.** Distributions of numbers of merozoites in individual schizonts. For each isolate, merozoites were counted in 100 schizonts (matured with chemical blocking of egress) as pooled counts from two different occasions (50 schizonts counted in each preparation). Moderate variation was seen, ranging from a mean of 16.5 for isolate 273 to 24.2 for isolate 293. **B.** Correlation between mean numbers of merozoites per schizont and multiplication rate assayed after day 153 of culture (Spearman's rho =0.46, P =0.20).

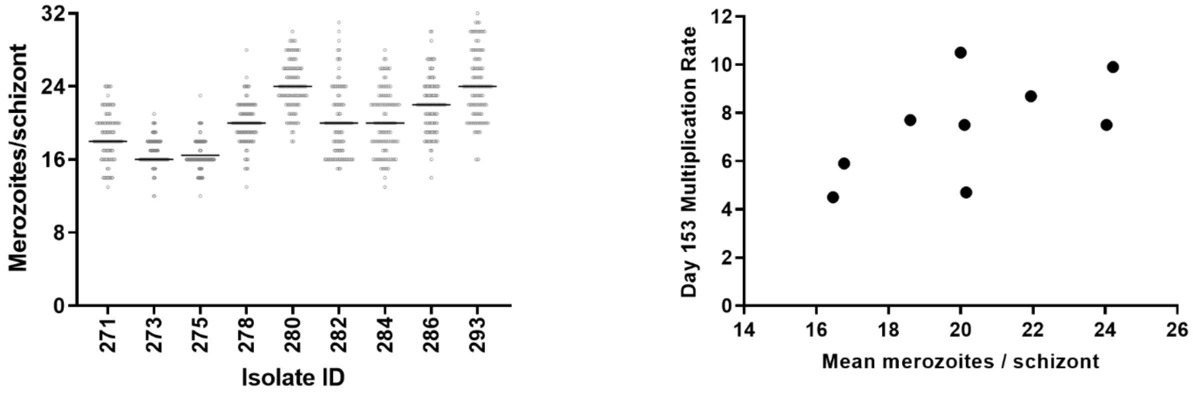
